# Whole Genome Sequencing based Identification of Human Tuberculosis caused by Animal-lineage *Mycobacterium orygis*

**DOI:** 10.1101/2023.02.27.530375

**Authors:** Md Rashedul Islam, Meenu K. Sharma, Rupinder KhunKhun, Cary Shandro, Inna Sekirov, Gregory J. Tyrrell, Hafid Soualhine

## Abstract

A recently described member of the *Mycobacterium tuberculosis* complex (MTBC) is *Mycobacterium orygis,* which can cause disease primarily in animals but also in humans. Although *M. orygis* has been reported from different geographic regions around the world, due to a lack of proper identification techniques, the contribution of this emerging pathogen to the global burden of zoonotic tuberculosis is not fully understood. In the present work, we report single nucleotide polymorphisms (SNPs) analysis using whole genome sequencing (WGS) that can accurately identify *M. orygis* and differentiate it from other members of MTBC species. WGS-based SNPs analysis was performed for 61 isolates from different Provinces in Canada that were identified as *M. orygis*. A total of 56 *M. orygis* sequences from the public databases were also included in the analysis. Several unique SNPs in *gyrB, PPE55, Rv2042c, leuS, mmpL6,* and *mmpS6* genes were used to determine their effectiveness as genetic markers for the identification of *M. orygis*. To the best of our knowledge, five of these SNPs, *viz., gyrB*^277^ (A→G), *gyrB*^1478^ (T→C), *leuS*^1064^ (A→T), *mmpL6*^486^ (T→C), and *mmpS6*^334^ (C→G) are reported for the first time in this study. Our results also revealed several SNPs specific to other species within MTBC. The phylogenetic analysis shows that studied genomes were genetically diverse and clustered with *M. orygis* sequences of human and animal origin reported from different geographic locations. Therefore, the present study provides a new insight into the high confidence identification of *M. orygis* from MTBC species based on WGS data, which can be useful for reference and diagnostic laboratories.

---

**T**uberculosis is a severe and complex infectious disease caused by the *Mycobacterium tuberculosis* complex (MTBC) and remains a public health concern leading to the death of 1·6 million people per year worldwide, predominantly in low-income and middle income countries (1). *Mycobacterium orygis,* a species belonging to MTBC, was first described by van Ingen et al. (2). *M. orygis* is capable of causing infection in both animal and human hosts. This species has received considerable interest in recent years, and has been reported to be isolated from dairy cattle and captive monkeys (3), captive wild animals (4), deer (5), rhinoceros (6), black buck (5), and humans (2, 7, 8). *M. orygis* infection has been recognized as a zoonotic source of human tuberculosis (9). Moreover, in New Zealand, a presumptive transmission of *M. orygis* from human to animal was reported, with the original infection being mapped out from contacts with domestic animals in India (10). *M. orygis* is endemic in Southeast Asian countries including Bangladesh, India, Nepal, and Pakistan (3, 7, 10). Although tuberculosis incidence in Canada is relatively low (11), zoonotic tuberculosis is progressively being recognized as a significant menace to the public health. Hence, zoonotic tuberculosis could pose a significant challenge in controlling this disease and meeting global tuberculosis elimination goals (12).

Along with *M. orygis*, other phylogenetically related MTBC species are *M. tuberculosis*, *M. bovis* and its variant BCG, *M. africanum*, *M. caprae*, *M. pinnipedii*, *M. microti*, *M. canettii*, and members of animal_Jadapted clade A1 such as “Dassie” bacillus, Chimpanzee bacillus, *M. suricattae,* and *M. mungi* (13–15). To improve human and animal health surveillance, it is important to implement proper identification methods and analysis tools for quickly and accurately discriminating MTBC species. Probes hybridization-based assays have been used to differentiate the causative agents of tuberculosis, however, studies have reported a limitation of this method in differentiating *M. orygis* from *M. africanum* (16). The mutations in genes including *gyrB* and *Rv2042c*, and the regions of difference (RDs) deletion have been used to discriminate *M. orygis* from MTBC species, using PCR-based approaches (2, 3, 17). Since *M. orygis* strains share *gyrB*^1450^ (G→T) mutation with MTBC members including *M. africanum* and *M. pinnipedii*, they may have previously been mislabeled as *M. africanum* (3, 17, 18).

Whole genome sequencing (WGS) technologies are now increasingly being used in clinical and research laboratories to investigate tuberculosis surveillance, outbreak detection, antimicrobial resistance prediction, characterization and diversity of MTBC species (14, 19). One of the key challenges for adopting WGS for these applications is data analysis that requires bioinformatics support and data interpretation (20). Consequently, a number of analytical tools have been developed to detect pathogenic bacterial strains using WGS data, with single nucleotide polymorphisms (SNP)-based methods being the common ones used in public health laboratories (21–23). BioHansel, for example, performs high-resolution genotyping by detecting phylogenetically informative SNPs in WGS data (24).

In order to improve our ability to accurately identify *M. orygis* and other species within MTBC, whole genome analysis was performed in this investigation. We report the identification of 61 new *M. orygis* from Canada by WGS-based SNP analysis, and to validate the results, publicly available 56 *M. orygis* genomes were added to the analysis. We performed molecular markers characterization on these genomes that demonstrate a clear differentiation of *M. orygis* from members of other MTBC species. Furthermore, the newly sequenced *M. orygis* isolates were phylogenetically analyzed to determine their diversity and global distribution. This study may improve our understanding of this poorly monitored emerging zoonotic pathogen and address its burden in Canada and globally.

## MATERIALS AND METHODS

### Sample collection, and project background

Cultures from across Canada were received at the National Reference Centre for Mycobacteriology (NRCM), National Microbiology Laboratory, Public Health Agency of Canada, Winnipeg for *M. tuberculosis* testing between 2009 and 2022. The cultures were grown on mycobacteria growth indicator tube (MGIT) media and Middlebrook 7H10 plates using standard and aerobic growth conditions. At NRCM, the presence of the insertion sequence IS*6110* and region of difference RD9 was confirmed by RT-PCR analysis to detect MTBC and *M. tuberculosis*, respectively. Classical methods such as *gyrB*, RD1, RD4, RD7 and mycobacterial interspersed repetitive unit-variable number of tandem repeat (MIRU-VNTR) were also used for identification and genotyping purpose. These methods can distinguish between mostly isolated MTBC species, *M. tuberculosis, M. bovis* and its variant BCG, *M. caprae* and *M. africanum*, but they are unable to identify some rare species of the complex such as *M. orygis* and *M. pinnipedii*. Since 2018, our group started using routinely WGS technologies and BioHansel (24) program for identification and differentiation between MTBC species. BioHansel performs high-resolution genotyping of MTBC by detecting phylogenetically informative SNPs in WGS data. However, the SNPs in the *gyrB* gene (at positions 432, 513, 870, 1068, 1167, and 1207) currently incorporated in BioHansel cannot differentiate *M. orygis*, and the pipeline eventually identifies this species, with low confidence, as a member of an animal lineage of the MTBC (probable *M. tuberculosis* var. *orygis*) based on SNP typing (2). We then separated all isolates identified as animal lineage “probable *M. orygis*”, all isolates with same MIRU-VNTR profile and *M. africanum* or *M. pinnipedii* or *M. bovis*/BCG from the inventory and investigated in this study. Therefore, a total of 137 isolates were interrogated (Supplementary Data S1), from which 61 were identified to be *M. orygis*. Most of the isolates were from British Columbia ‘BC’ (n = 38), followed by Alberta ‘AB’ (n = 19), Manitoba ‘MB’ (n = 2), Nova Scotia ‘NS’ (n = 1), and Saskatchewan ‘SK’ (n = 1). The detailed information (*e.g.,* source Province, specimen type, and identification) on *M. orygis* isolates from this study is depicted in Table 1.

**TABLE 1.**
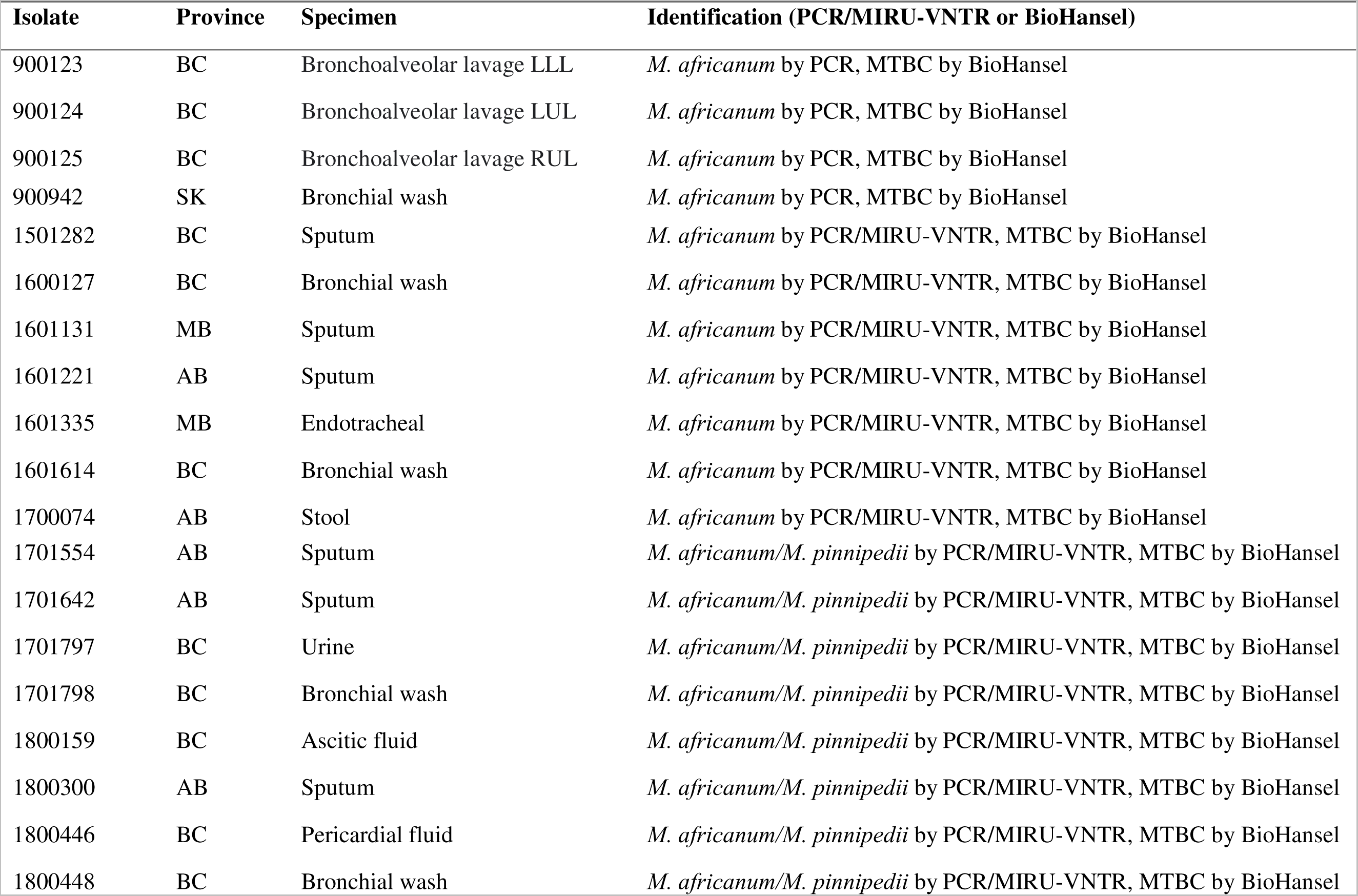
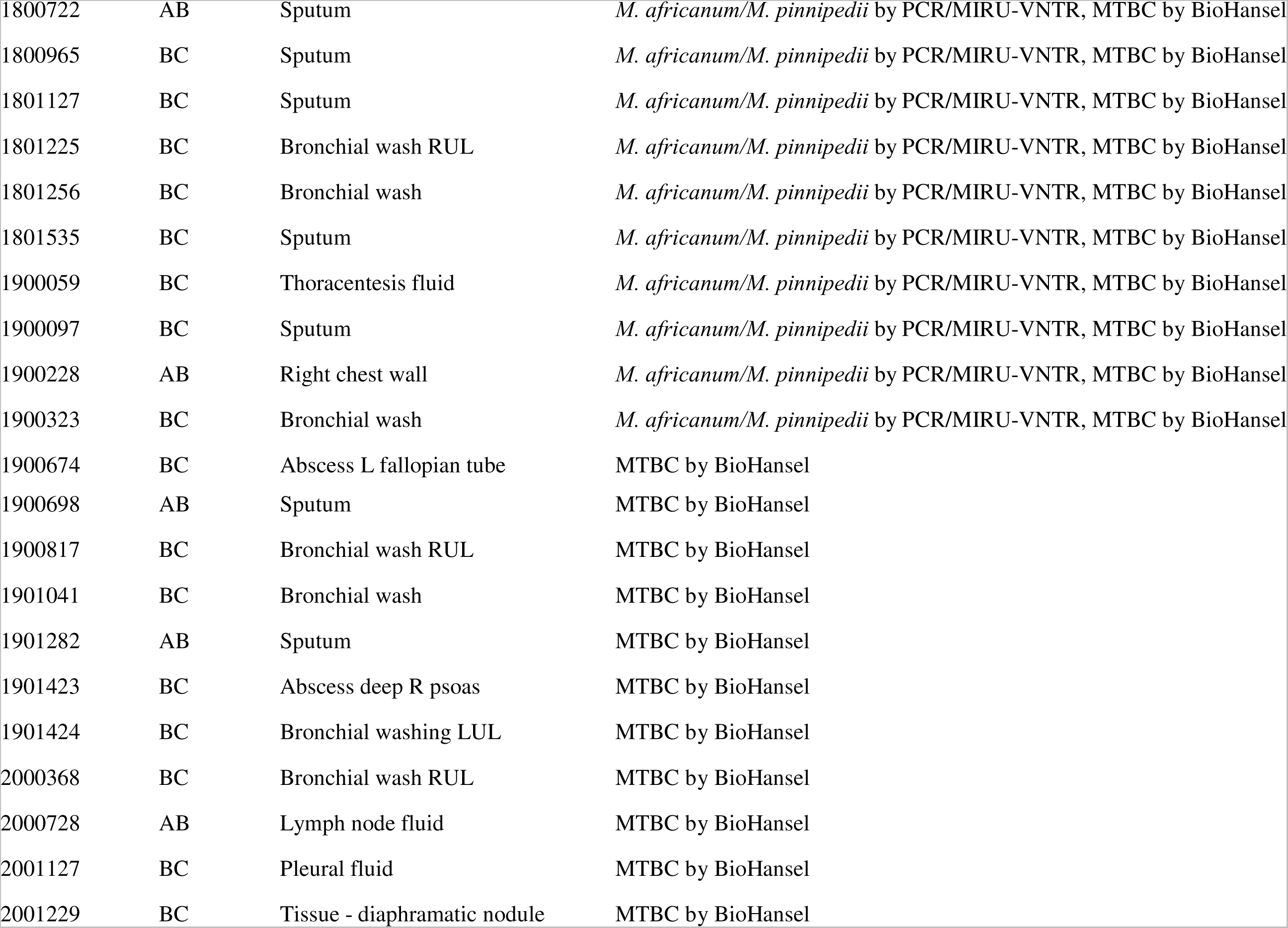
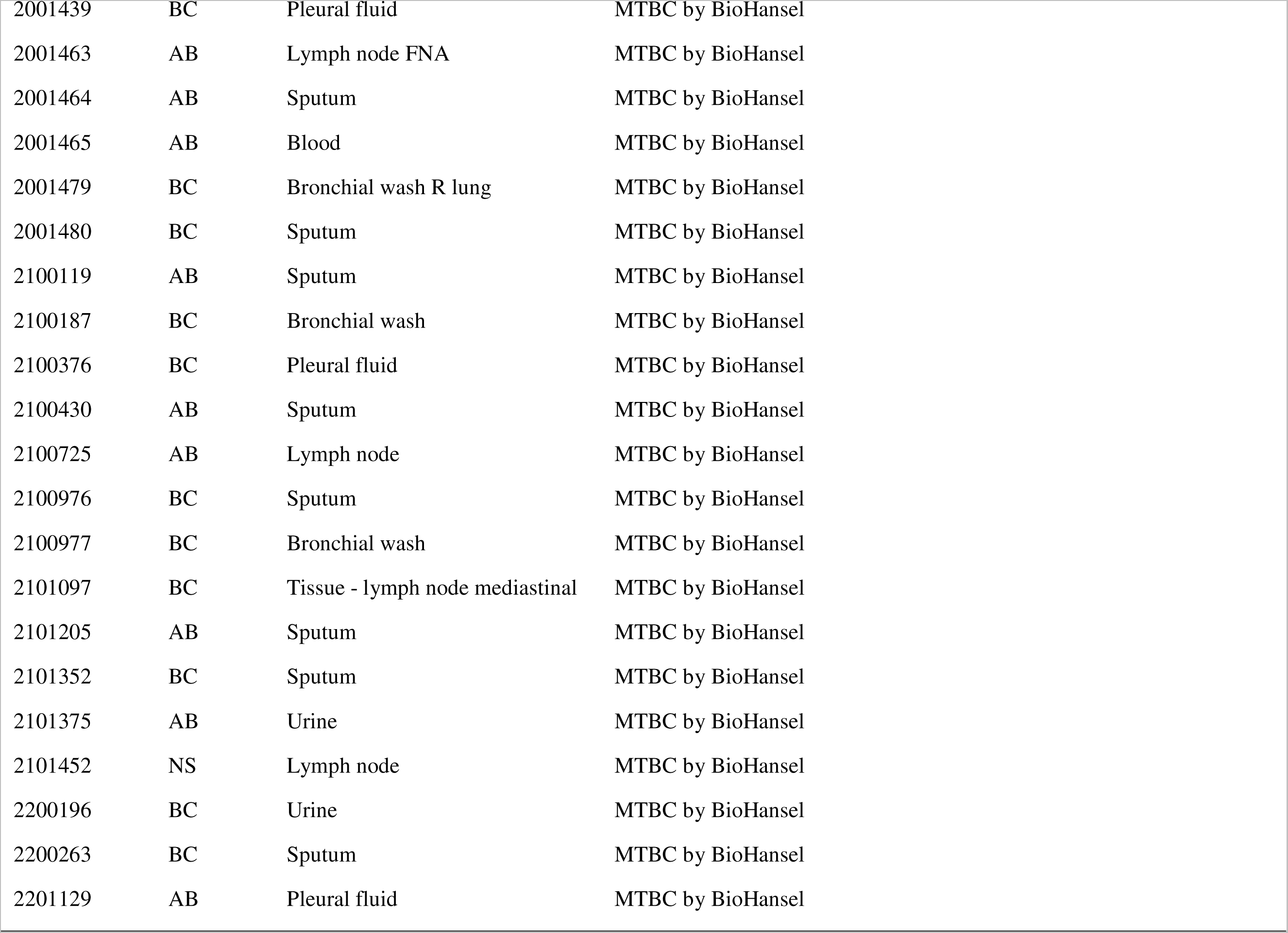
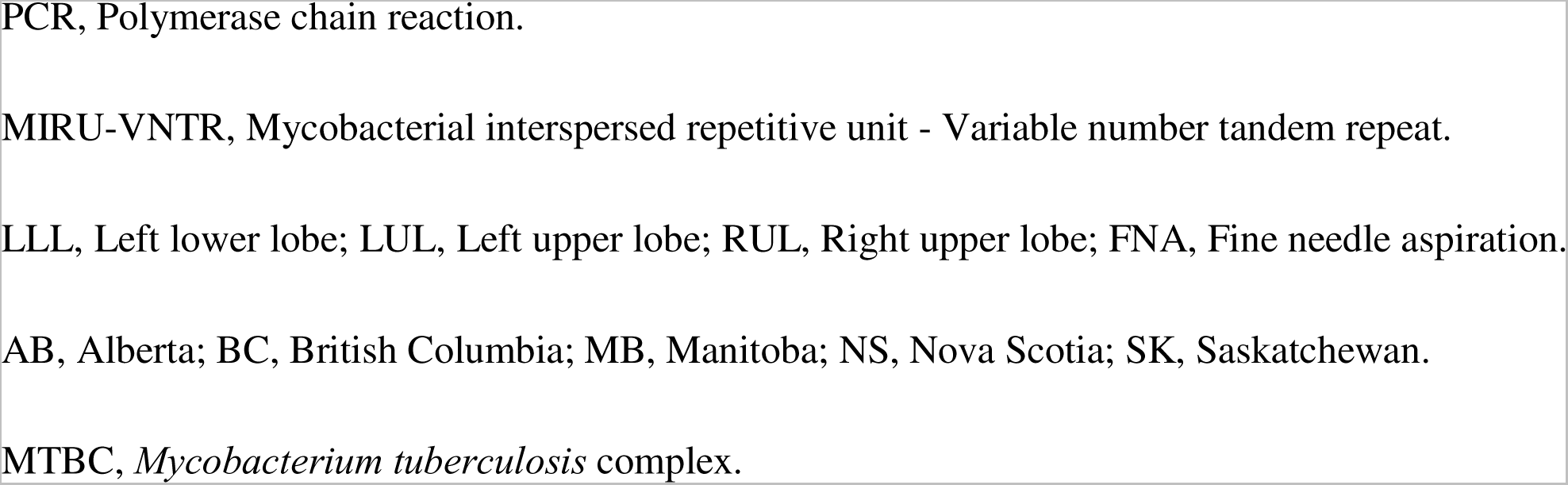
Newly identified 61 *M. orygis* isolates from this study.

### Whole genome sequencing (WGS). (i) DNA extraction, sequencing and data processing

Genomic DNA was extracted from the isolates using the InstaGene^TM^ Matrix comprising 6% (*w/v*) Chelex resin for PCR-ready DNA purification (Bio-Rad #7326030; California, United States). DNA quality was checked by measuring the *A260/280* ratio using NanoDrop, and quantified fluorometrically using the Qubit 3.0 (Thermofisher, Massachusetts, United States). The sequencing library was prepared using the Illumina DNA Prep Kit (Illumina, California, United States), according to the manufacturer’s protocol. Paired-end sequencing was conducted on an Illumina MiSeq platform (Illumina Inc.) using 300-cycle MiSeq Reagent Kit v2, 500-cycle MiSeq Reagent Kit v2 or 600-cycle MiSeq Reagent Kit v3 (Illumina, California, United States), creating 2 × 150, 2 × 250 or 2 × 300 bp paired-end reads, respectively.

Raw sequencing reads were processed in the IRIDA platform (25) (http://irida.ca) using Shovill for assembly (26), Prokka for annotation (27), and QUAST for assembly assessment (28). Sequencing Report workflow v.2.3, where FastQC (29) and SMALT (https://www.sanger.ac.uk/tool/smalt-0/) are the two main tools, was implemented in the Galaxy platform (30) for all studied isolates by aligning the reads against the reference genome of *M. tuberculosis* H37Rv (accession # NC_000962.3). Only sequences with at least 60% of mapped reads and a >28x average coverage were filtered for further analysis. While 54 out of 61 studied *M. orygis* sequences had mapped reads of over 80%, the remaining seven sequences were found to have 60-80% of mapped reads, and only one sequence (2100725) had average coverage below 30x (28.85).

### (ii) Genomic comparison of the new sequences with public database

The newly sequenced genomes were compared with 56 publicly available *M. orygis* genomes reported from around the world, downloaded from the National Center for Biotechnology Information (NCBI) GenBank (31), the NCBI Sequence Read Archive (32), and the European Nucleotide Archive (33). The list of reference *M. orygis* genomes with their accession numbers, collection centres, country of origin, host species and specimen source, is presented in Supplementary Data S2A. Aside from *M. orygis*, a total of 190 previously published genome sequences (complete/draft/SRA) representing all MTBC species and lineages (L1 to L9), including its animal lineages, were also downloaded from the above repositories and added to the analysis (Supplementary Data S2B).

### Gene selection and R analysis: Unique SNPs for *M. orygis* and other MTBC species/lineages

To develop a simple, quick, effective and yet powerful identification approach, a set of six genes, *viz., gyrB, PPE55, Rv2042c, leuS, mmpL6,* and *mmpS6* were selected for SNPs analysis, and these genes were chosen based on the previous publications (2, 17, 34). The sequence for *gyrB, PPE55, Rv2042c* and *leuS* genes were obtained from H37Rv (accession # NC_00962.3), and *mmpL6* and *mmpS6* gene sequences were recovered from Mtb-specific deletion 1 region ‘TbD1’ (accession # AJ426486.1). The numerical positions for the new and known SNPs in the *gyrB, Rv2042c, PPE55,* and *leuS* genes are relative to *M. tuberculosis* H37Rv reference genome (accession # NC_000962.3, (35), while the SNPs positions within *mmpL6* and *mmpS6* genes are provided following TbD1 region (AJ426486.1, (36)). All identified SNPs within the six genes are numbered according to gene position (5_J to 3_J), and the complementary sequences were used for genes *Rv2042c* and *PPE55* (Table 2).

**TABLE 2.**
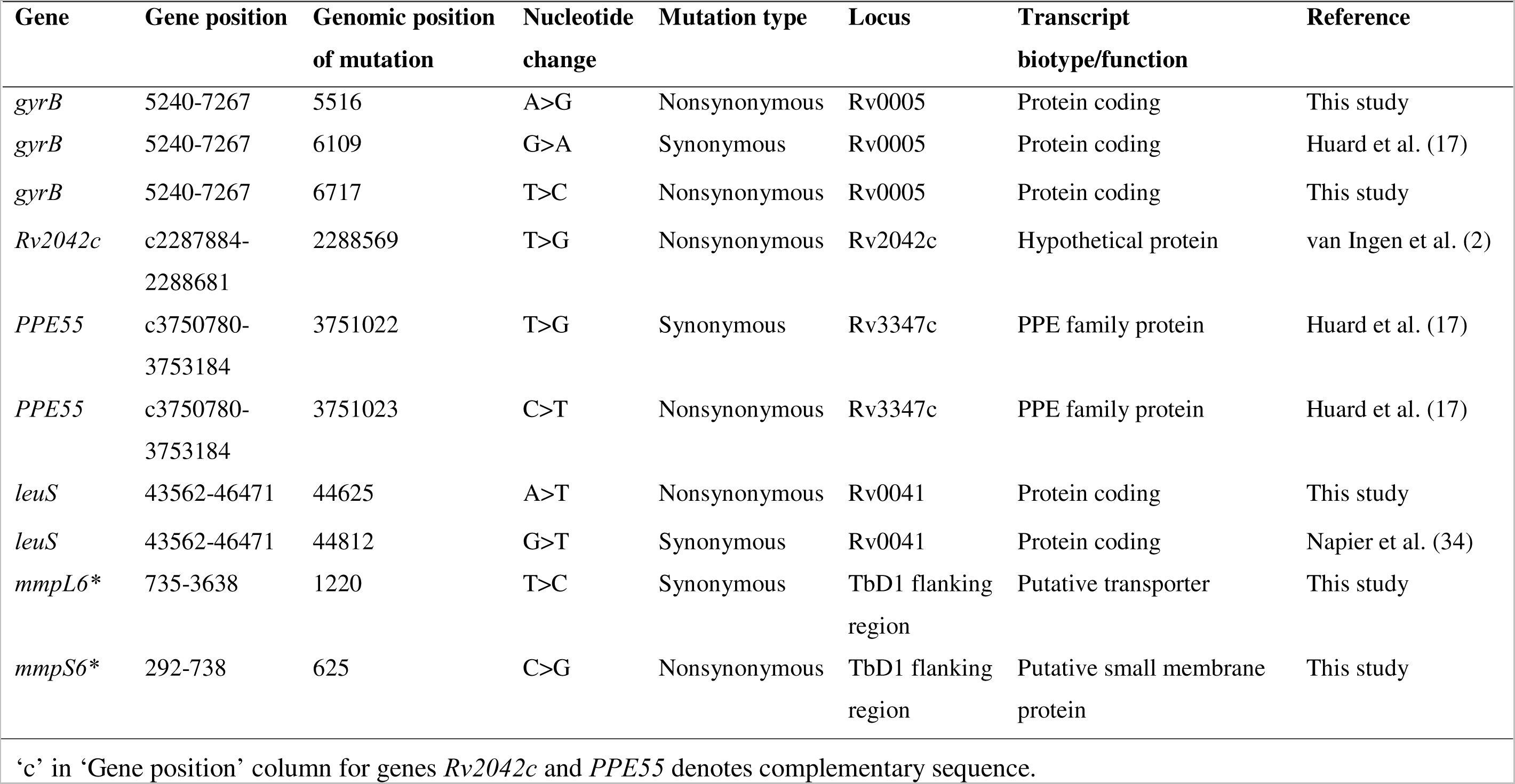
Signature single nucleotide polymorphisms (SNPs) in *M. orygis*, as referenced to H37Rv (NC_000962.3) and Mtb-specific *deletion* 1 region ‘TbD1’ (AJ426486.1*)

To extract the above-mentioned six gene sequences from *M. orygis* and MTBC species/lineage genomes, a high-throughput analysis was performed using our custom in-house-developed R-scripts in RStudio v.2021.9.2.382 (37), which used Basic Local Alignment Search Tool (BLAST) (38), with the e-value cutoff option set to 10e-100. The scripts interrogated the assembled genomes using a reference gene to identify and extract the sequence corresponding to the SNPs of interest. NCBI BLAST search was also done for all genes, with any unique sequences investigated. In order to identify *M. orygis* or other MTBC species-specific new and known SNPs, the extracted gene sequences were aligned using MUSCLE (39), and sequence variations of the unique SNPs were detected using MEGA-X v.10.0.5 (40) and Geneious Prime 2022 (Geneious Prime 2022.0.2, https://www.geneious.com).

### The SNPs-based phylogenetic reconstruction of *M. orygis*

To compare *M. orygis* sequences from the present study with those available in the public repositories, a whole genome phylogeny was analyzed using the single nucleotide variant phylogenomics (SNVPhyl) pipeline v.1.2.3 in IRIDA (23). The snvAlignment file obtained from SNVPhyl was then used to construct a maximum likelihood tree using RAxML v.8.2.12 with the GTRGAMMA model and 100 rapid bootstrap replicates (41). The output phylogenetic tree was visualized using the online tool iTOL (Interactive Tree Of Life) v.6 (42). *M. tuberculosis* H37Rv (NC_00962.3) was chosen as a reference genome with parameters: minimum SNV coverage of 15, SNV abundance ratio of 0.75, and minimum mapping quality of 30. The SNV density filtering was enabled to remove high-density SNV regions that could suggest possible recombination. The identified SNVs were used in the construction of a phylogeny by calculating genetic distance between isolates by a generalized time reversible (GTR) model with PhyML 3.0 (43).

### Data availability

The whole genome sequences of *M. orygis* isolates from this project have been deposited in GenBank and are available through the SRA under BioProject no. PRJNA934340 with the submission ID SUB12537725. Accession numbers for all publicly available genomes used in this study are listed in Supplementary Data S2.

## RESULTS

### Data collection and genome analysis

In the present study, a total of 61 *M. orygis* isolates were identified retrospectively from cultures submitted by five different Provinces in Canada over a period of 14 years (2009-2022), with more than 90% from British Columbia and Alberta. The cultures were recovered from pulmonary (65%) and extra-pulmonary samples. Forty-five *M. orygis* cases were female patients (74%), 14 were male patients (23%) and two cases did not provide gender on the requisition. The PCR, MIRU-VNTR testing, BioHansel and Mykrobe Predictor v.0.7.0 (44) analyses identified the isolates as *M. africanum* or *M. africanum/M. pinnipedii* or MTBC animal lineages (Table 1). We then performed WGS for all 61 isolates and determined their identification as *M. orygis* using SNPs analysis. The sequencing data were augmented with a total of 56 publicly available *M. orygis* genome sequences to validate the identification and their discrimination from other MTBC species. Of these 56 sequences, 35 were reported from India, followed by eight from The United States, five from Norway, four from Switzerland, two from The United Kingdom, one from The Netherlands, and one from Canada (Supplementary Data S2A).

### Whole genome-based SNPs analysis

We performed the WGS-based SNPs analysis for identification and genetic differentiation of *M. orygis* from other MTBC members. Several new and known SNPs in *gyrB* (2028 bp)*, PPE55* (2405 bp)*, Rv2042c* (798 bp)*, leuS* (2910 bp)*, mmpL6* (2904 bp), and *mmpS6* (447 bp) genes were evaluated to ascertain their effectiveness as molecular markers for *M. orygis*. By gene sequence analysis of multiple sequence alignment from 117 *M. orygis* genomes (61 studied and 56 public repositories) and 190 sequences of MTBC species and lineages, we identified 10 unique SNPs across six selected genes that can identify and discriminate *M. orygis* from all members of MTBC. These *M. orygis* specific SNPs along with their corresponding genomic positions, mutation type, locus, and function are listed in Table 2.

A multiple alignment of the *gyrB* gene showed the presence of two novel nonsynonymous mutations at positions 277 (A→G) and 1478 (T→C), along with a known synonymous mutation at position 870 (G→A) (Table 2, Table 3). Within *PPE55* gene, our study identified two *M. orygis* specific SNPs at positions 2162 (C→T) and 2163 (T→G). The SNPs analysis also identified a unique SNP in *Rv2042c* gene at position 113 (T→G). In addition, we found two *M. orygis* specific SNPs within *leuS* gene; one of which was a novel SNP (*leuS*^1064^ A→T) and associated with synonymous mutation, and another one (*leuS*^1251^ G→T) was previously reported. The sequence of the TbD1 region (AJ426486.1) from the ancestral *M. tuberculosis* contains genes *mmpS6* and *mmpL6*. Two more novel SNPs were detected in this region and were found to be associated with a synonymous (*mmpL6*^486^ T→C) and a nonsynonymous (*mmpS6*^334^ C→G) mutation (Table 2, Table 3). The genes *mmpL6* and *mmpS6* encode membrane proteins belonging to a large and a small family, respectively. The above 10 unique SNPs situated within a set of six genes were identified in all 117 *M. orygis* strains screened in the present study.

**TABLE 3.**
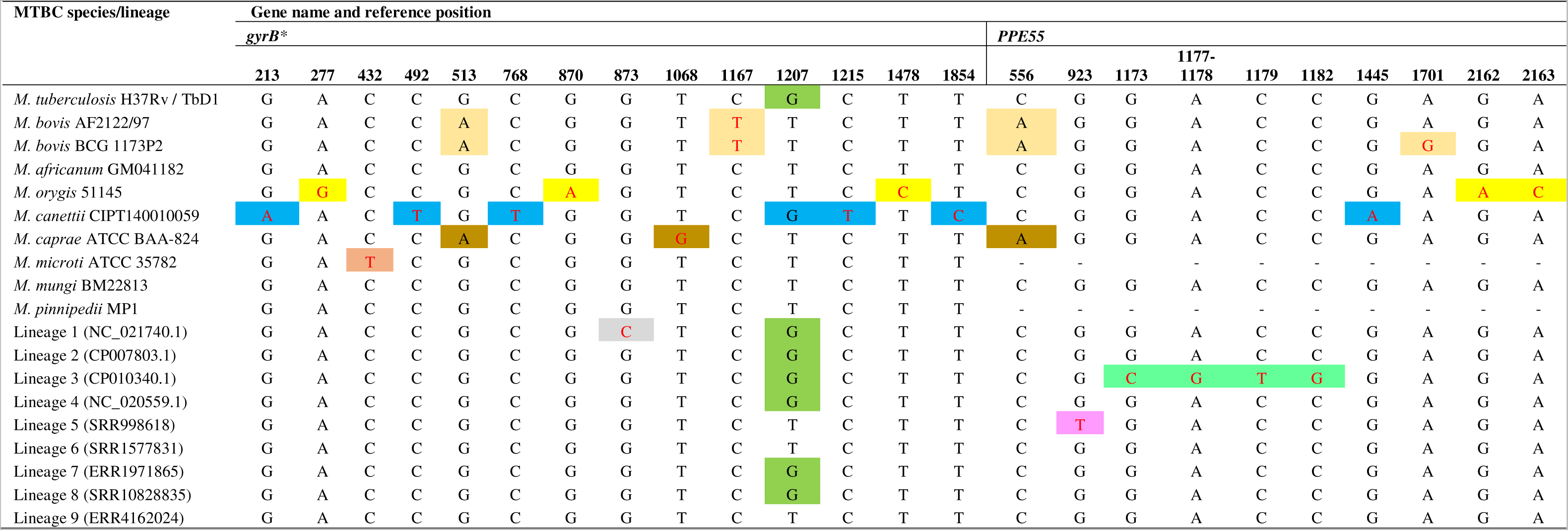
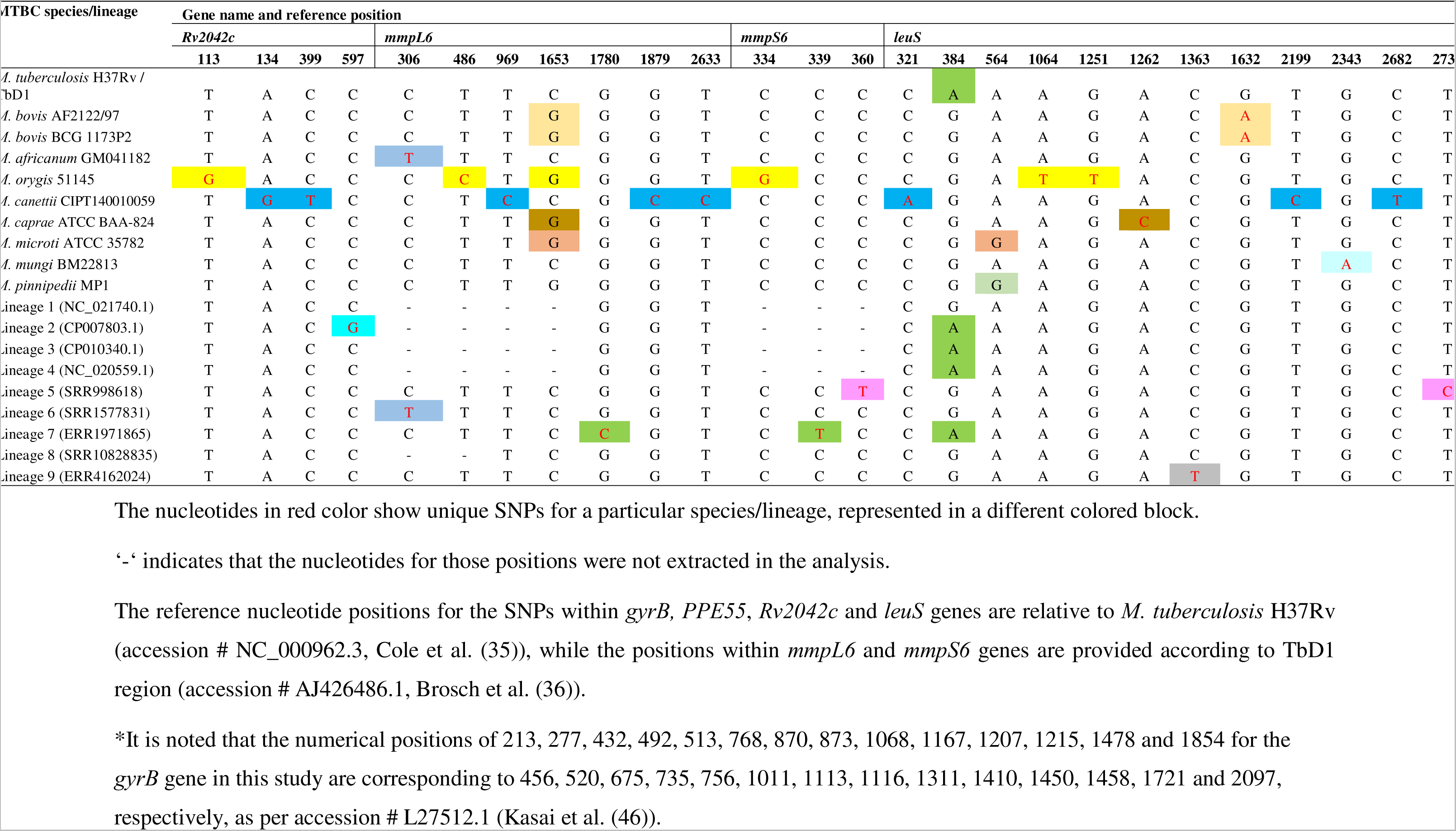
Summary of single nucleotide polymorphisms (SNPs) analysis for *M. orygis* differentiation from other MTBC species and lineages. Fourteen SNPs on *gyrB,* eleven SNPs on *PPE55*, four SNPs on *Rv2042c*, seven SNPs on *mmpL6*, three SNPs on *mmpS6*, and twelve SNPs on *leuS* genes are shown.

In order to define the discriminatory power of the WGS-based SNPs analysis in identifying all MTBC species/lineages, the above 117 *M. orygis* (Table 1, Supplementary Data S2A) and 190 MTBC (Supplementary Data S2B) genome sequences were also analyzed, and results of representative sequence from each species and lineage are shown in Table 3. Apart from *M. orygis* specific SNPs, we identified some new and previously reported SNPs that are unique for all MTBC members and lineages. These include 11 SNPs in *gyrB*, 10 SNPs in *leuS*, nine SNPs in *PPE55*, six SNPs in *mmpL6*, three SNPs in *Rv2042c*, and two SNPs in *mmpS6* genes.

In our study, we detected a series of SNPs specific for *M. canettii* in all six selected genes, including *gyrB* (213G→A, 492C→T, 7681C→T, 1215C→T and 1854T→C), *PPE55* (1445G→A), *Rv2042c* (134A→G), *mmpL6* (1879G→C), and *leuS* (321C→A and 2199T→C). Species-specific polymorphisms were also found for *M. caprae* (*gyrB*^1068^ T→G and *leuS*^1262^ A→C), *M. microti* (*gyrB*^432^ C→T) and *M. mungi* (*leuS*^2343^ G→A). Interestingly, one SNP identified in *leuS* gene (564A→G) of *M. microti* was overlapped in *M. pinnipedii* strains. The gene sequence analysis identified a SNP (*mmpL6*^306^ C→T) that can differentiate *M. africanum* lineage 6 from other MTBC members and lineage. While the SNPs in *gyrB* gene at position 1167 (C→T) and in *leuS* at position 1632 (G→A) were found to discriminate *M. bovis* and *M. bovis* BCG from all others, the SNP *PPE55*^1701^ A→G could even separate *M. bovis* from BCG. Furthermore, *M. bovis* and BCG shared SNPs *gyrB*^513^ G→A and *PPE55*^556^ C→A with *M. caprae* strains (Table 3). In addition to the above, we detected SNP markers in the selected six genes that differentiated the strains belonging to different lineages from other MTBC members examined in this study.

### Single nucleotide polymorphisms-based phylogenetic inference of *M. orygis*

A whole genome SNP-based phylogenetic tree was built to determine the genetic similarity of the newly sequenced *M. orygis* isolates in Canada with *M. orygis* sequences from the public repositories (Fig. 1). The SNV distance matrix generated from the whole genome alignments lists the pairwise SNV distances between every sequence (Supplementary Data S3). The genome sequences from this study clustered with *M. orygis* sequences reported from all five geographic locations.

**FIG 1.**
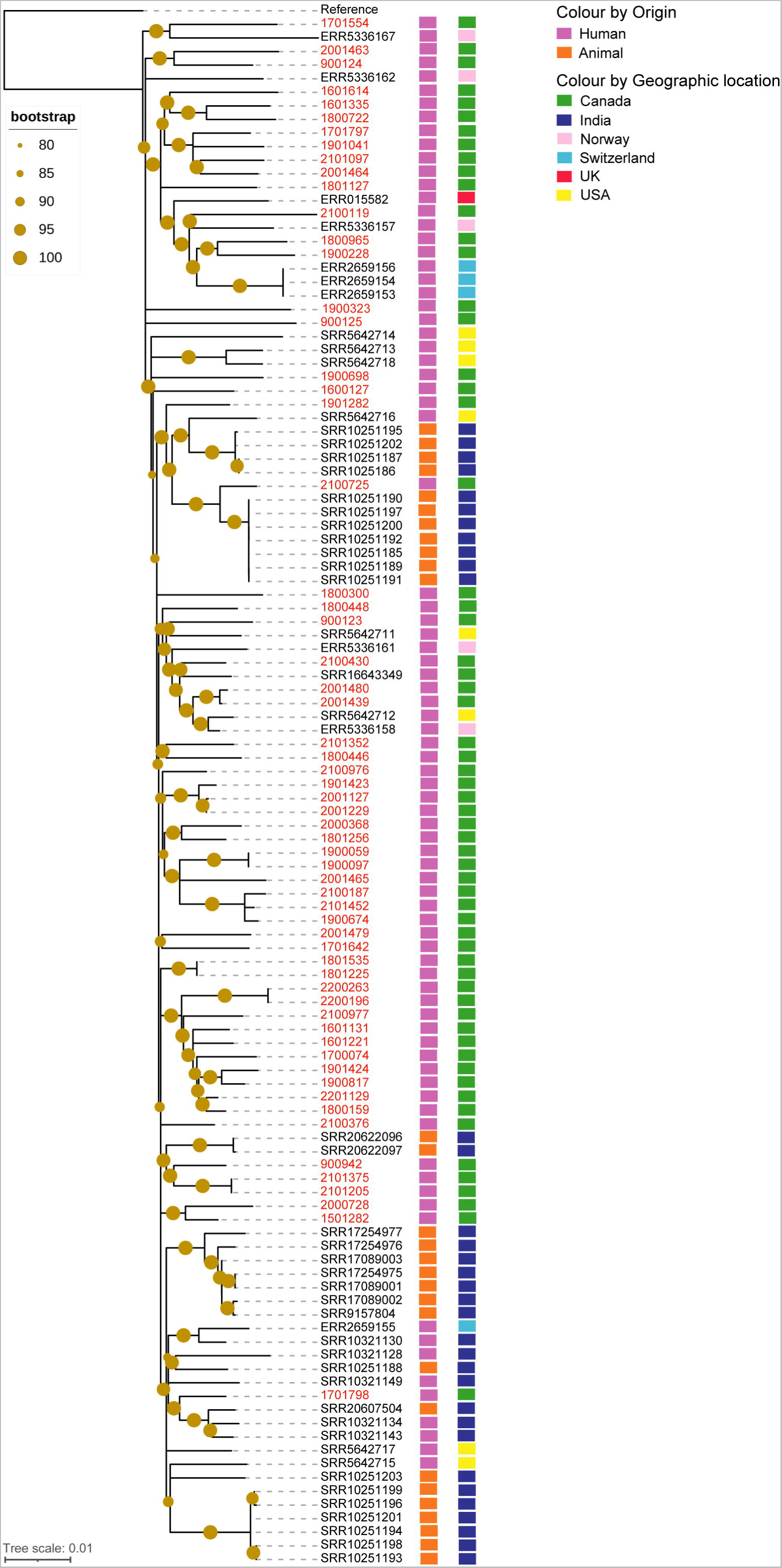
Phylogenetic tree of *M. orygis* based on single nucleotide variants (SNVs) analysis. The whole genome phylogeny was analyzed using SNVPhyl pipeline v.1.2.3, and the snvAlignment file was used to infer the tree using RAxML v.8.2.12 with the GTRGAMMA model and 100 rapid bootstrap replicates. The output tree was visualized using iTOL v.6. A maximum likelihood tree of 61 newly sequenced *M. orygis* isolates from this study (labelled in red), and the *M. orygis* genomes collected from public repositories. The left bar adjacent to the tree nodes shows the source of *M. orygis* (human in purple, and animal in orange), and the right bar denotes shows the geographic location of *M. orygis* (Canada in green, India in blue, Norway in pink, Switzerland in cyan, UK in red, and USA in yellow). *M. tuberculosis* H37Rv (NC_000962.3) used as a reference in this analysis. The filled circles on nodes represent a bootstrap value between 80 and 100. The studied isolates mostly clustered with *M. orygis* of human origin recovered from different geographic regions. The isolate 2100725 showed a good phylogenetic relation (pairwise distances of 34 SNVs) with seven *M. orygis* of animal origin from India.

*M. orygis* sequences from different samples of the same patient clustered together in the SNVPhyl tree, as expected. For example, BC isolate 1801225 clustered with 1801535, 1900059 with 1900097, 2001127 with 2001229, 2001439 with 2001480 and 2200196 with 2200263, and AB isolate 2101205 with 2101375 appeared to have a SNVs distance of 0-5. The isolates 2001439 and 2001480 also clustered (29-52 SNVs distance) with two *M. orygis* sequences from Norway (ERR5336158) and The United States (SRR5642712), and one previously reported *M. orygis* sequence 51145 (SRR16643349) from Quebec Canada. Similarly, 11 *M. orygis* sequences from the present study formed a large cluster with three sequences (ERR2659153, ERR2659154, ERR2659156) from Switzerland, one sequence (ERR5336157) from Norway, and another sequence (ERR015582) reported from The United Kingdom, with two sequences (1800965 from BC and 1900228 from AB) showing a distance of 92-96 SNVs with the *M. orygis* sequences from Switzerland (Fig. 1, Supplementary Data S3).

The studied isolates 1901282 and 2100725 from AB clustered with 11 *M. orygis* sequences of animal origin from India and one sequence of human origin from The United States with a pairwise SNV distance of 34-88. Interestingly, the isolate 2100725 clustered with seven of these animal originated sequences (SRR10251185, SRR10251189-92, SRR10251197, and SRR10251200) by 34 SNV distances. However, the SNV numbers of 34-88 in MTBC are quite a large distance and are not indicative of spillover or direction of spillover.

## DISCUSSION

The present study is the first to report a greater number of *M. orygis* from a single geographic area. *M. orygis* is a causative agent of tuberculosis in both animal and human hosts and it was first described in 2012 (2), and later by others (3–5, 7, 8). The MTBC species *M. bovis* has long been believed to be the only agent that causes zoonotic tuberculosis, however, recovering a larger number of *M. orygis* from both animals and humans in recent years from different areas of the world highlights the need for considering this bacterium as a zoonotic pathogen (45).

Since tuberculosis cases caused by *M. orygis* are often identified as MTBC or misidentified and published as *M. africanum* or *M. bovis*, the actual number of infections associated with this bacterium may have been underreported (3, 16, 46). From our culture collection, the investigation showed that since 2009, *M. orygis* was misidentified as *M. africanum*. Rahim et al. (18) initially reported *M. africanum* from four dairy cows that were later identified to be *M. orygis* by refined analysis (3). A part of the confusion towards this misidentification is that *M. orygis* shares the *gyrB*^1207^ (G→T) (*gyrB*^1450^ as per accession no. L27512.1, (47)) mutations with *M. africanum*, *M. bovis, M. microti*, *M. caprae, M. mungi*, and *M. pinnipedii* ((17), Table 3). Moreover, MTBC is genetically highly clonal, and thus without proper identification tools and analysis approaches the species differentiation could be challenging.

Since currently available laboratory tests are struggling to differentiate animal lineages, in the present investigation we used WGS-based SNPs analysis targeting *gyrB, PPE55, Rv2042c, leuS*, *mmpL6,* and *mmpS6* genes that can accurately identify *M. orygis* and unambiguously differentiate it from all members of MTBC species. The *gyrB* gene encodes for the β-subunit of the DNA gyrase and has been used as a molecular marker for identification of MTBC members. The discriminatory power of polymorphisms in *gyrB* gene in identifying *M. orygis* has been evaluated in previous studies (3, 17). While these authors described only one unique SNP (*gyrB*^870^ G→A) (*gyrB*^1113^ according to accession no. L27512.1), we detected two more novel and useful genetic markers within *gyrB* gene (*gyrB*^277^ A→G, and *gyrB*^1478^ T→C) in the screened *M. orygis* genomes. The *PPE55* (Rv3347c) gene is specific to MTBC species and play a major role in host-pathogen interactions. Huard et al. (17) earlier reported two *M. orygis* specific SNPs in *PPE55* gene (*PPE55*^2162^ C→T, and *PPE55*^2163^ T→G), that was also revealed in our study. The results suggest that these SNPs could be used as genetic markers for the identification of *M. orygis*. We also evaluated *Rv2042c* gene for possible unique SNPs to be used as a molecular marker to identify *M. orygis*. In accordance with van Ingen et al. (2), our SNPs analysis identified a nonsynonymous mutation in the 38th codon of *Rv2042c* gene at position 113 (T→G). In mycobacterial species, *leuS* gene encodes for L-leucyl-tRNA synthetase, which is involved in translation. In this study, in addition to the SNP *leuS*^1251^ (G→T) described by Napier et al. (34), we showed that another novel SNP *leuS*^1064^ (A→T) associated with nonsynonymous mutation is also present in *M. orygis* strains (Table 2, Table 3). Mtb-specific deletion 1 region ‘TbD1’ region comprises the *mmpL6* and *mmpS6* genes, which encode for mycobacterial membrane protein families. In modern *M. tuberculosis* strains, the *mmpS6* gene is fully deleted and the *mmpL6* gene is trimmed (48). Our result shows two SNP *mmpL6*^486^ T→C, and *mmpS6*^334^ C→G, as two novel and distinct genetic markers for the identification of *M. orygis*.

We included previously described *M. orygis* specific polymorphisms in the WGS-based SNPs analysis, and they were reconfirmed in a larger number (61 studied and 56 from public repositories) of *M. orygis* genomes tested in this work. Thus, the results suggest WGS-based SNPs analysis is a useful tool to rapidly identify *M. orygis*, and to clearly differentiate this emerging pathogen from all MTBC species and lineages. To the best of our knowledge, the unique SNPs *gyrB*^277^ (A→G), *gyrB*^1478^ (T→C), *leuS*^1064^ (A→T), *mmpL6*^486^ (T→C), and *mmpS6*^334^ (C→G) are the first to be reported in the present study.

WGS-based SNPs analysis has been employed for the identification of MTBC species (9), and this approach has also become pertinent to MTBC genotyping (36, 49). In this study, we further identified the species-specific polymorphisms for members of MTBC species (excluding *M. orygis* here, as discussed above) and lineages, and results from *gyrB* gene confirm the discrimination of several MTBC members including *M. tuberculosis*, *M. bovis, M. canettii*, *M. microti,* and *M. caprae*, (Table 3). These results are in agreement with Huard et al. (17), who reported the identification of the above species (along with *M. orygis*) using a PCR-based SNPs analysis. Our study supports that many genomic characteristics could be shared between *M. bovis* and *M. caprae* strains, for example, the *gyrB* mutation at position 513, and *PPE55* mutation at position 556 (17).

Furthermore, we detected unique polymorphisms for lineage 1 in *gyrB*^873^ gene (G→C); for lineage 2 in *Rv2042c*^597^ (C→G) gene; for lineage 3 in *PPE55* gene at positions 1173 (G→C), 1177-1178 (A→G), 1179 (C→T) and 1182 (C→G); for lineage 5 in *PPE55*^923^ (G→T), *mmpS6*^360^ (C→T) and *leuS*^2736^ (T→C) genes; for lineage 7 in *mmpL6*^1780^ (G→C) and *mmpS6*^339^ (C→T) genes; and for lineage 9 in *leuS*^1363^ (C→T) gene. The reason for not being able to extract some SNPs in *mmpL6* and *mmpS6* genes for *M. tuberculosis* H37Rv and lineages 1-4 in Table 3 is that these genes are partly or fully deleted from the modern *M. tuberculosis*, as discussed above. Our results indicate that WGS-based SNPs analysis could be successfully used to distinguish all members of MTBC species and lineages. Since we tested a limited number of sequences due to availability and data quality, particularly for *M. pinnipedii, M. caprae,* and *M. mungi*, further investigation with a large dataset would be useful for their specific identification with high confidence.

The use of whole genome SNPs-based phylogenetic tree allowed us to inspect the genetic relation of *M. orygis* recovered from Canada and those reported from other geographic regions. The *M. orygis* isolates of human origin from this study were found to be distributed across the phylogeny (Fig. 1). Phylogenetic analysis of *M. orygis* genomes from the same patient on the same episode (two isolates from each of the six patients) showed a difference of 0-5 SNV.

We sought to determine whether *M. orygis* identified in this study showed a close phylogenetic relation with animal strains. The result shows that the isolate 2100725 from an AB patient clustered (34 SNVs apart) with several *M. orygis* sequences of animal origin reported from India (Fig. 1, Supplementary Data S3). This result may be indicative of the adaptation of animal origin *M. orygis* strains to a human host. The zoonotic or zooanthroponosis potentials of *M. orygis* has been discussed in a previous work (45).

Furthermore, two isolates from BC were related (5-39 SNVs apart) to *M. orygis* sequences of human origin reported from Norway (ERR5336158, (8) and The United States (SRR5642712, (7), and segregated by 49-52 SNVs from previously published Canadian strain 51145 (50). Thus, the newly sequenced *M. orygis* from Canada phylogenetically clustered with *M. orygis* sequences reported from different geographic locations, placing it within the global context.

In conclusion, the WGS analysis in the present study evaluated 10 novel and known unique SNPs within a set of six genes that could be used as molecular genetic markers to accurately identify *M. orygis* and unambiguously discriminate it from all members of MTBC species and lineages. As WGS technologies are increasingly being used by the healthcare systems, our approach will be helpful to diagnosis and surveillance of *M. orygis* associated tuberculosis, and optimizing the clinical management of this disease. The analysis of *M. orygis* sequences will improve our understanding of molecular characteristics and phylogenetic diversity of this emerging pathogen and its implication as a zoonosis. The ever-increasing evidence of *M. orygis* linked endemicity, and the identification of a greater number of *M. orygis* from animals and humans around the world highlight the urgency for a multi-sectoral collaboration linking the clinical and veterinary sectors towards One Health approach. The origin, epidemiology and transmission dynamics of *M. orygis* within Canada is currently under investigation.

## ACKNOWLEDGMENTS

The authors would like to thank Darrell Johnstone, Debra Janella and members of the National Reference Centre for Mycobacteriology (NRCM) and DNA Sequencing Core Services for their technical assistance with this project. The authors also acknowledge Walter Demczuk for help with R-scripting, and Aaron Petkau for valuable suggestions with phylogenetic analysis. This work is supported under the Genomics Research and Development Initiative (GRDI) program funded by Public Health Agency of Canada.

## ETHICS STATEMENT

This study was exempted from Health Canada and Public Health Agency of Canada Research Ethics Board (REB) since no clinical samples or human biological materials were used for this study.

